# Characterization of the endocrine pancreas in Type 1 Diabetes: islet size is maintained but islet number is markedly reduced

**DOI:** 10.1101/480509

**Authors:** Peter Seiron, Anna Wiberg, Lars Krogvold, Frode Lars Jahnsen, Knut Dahl-Jørgensen, Oskar Skog, Olle Korsgren

## Abstract

Insulin deficiency in type 1 diabetes (T1D) is generally considered a consequence of specific beta-cell loss. Since healthy pancreatic islets consist of ~65% beta cells, this would lead to reduced islet size if the beta cells are not replaced by other cells or tissue. The number of islets per pancreas volume (islet density) would not be affected.

In this study, we compared the islet density, size, and size distribution in subjects with recent-onset or long-standing T1D, with that in matched non-diabetic subjects. Results show that subjects with T1D, regardless of disease duration, had a dramatically reduced islet number per mm^2^, while the islet size was similar in all groups. Insulin-negative islets in T1D subjects were dominated by glucagon-positive cells that frequently had lost the alpha-cell transcription factor ARX while instead expressing PDX1, normally expressed in beta cells.

Based on our findings, we propose that failure during childhood to establish a sufficient islet number to reach the beta-cell mass needed to cope with episodes of increased insulin demand contributes to T1D susceptibility. Exhaustion induced by relative lack of beta cells could then potentially drive beta-cell dedifferentiation to alpha-cells, explaining the preserved islet size observed in T1D compared to controls.

## Introduction

The number of islets in the human pancreas is about 1-2 million with an absolute majority of islets with a diameter below 50 *µ*m. However, the islets with largest contribution to the total endocrine volume have a diameter of about 120-150 *µ*m. A tremendous expansion of the total islet volume occurs during childhood (1). If this expansion of the islet mass mainly occurs from a so far not identified pancreatic endocrine stem cell or via cell replication remains unknown (2; 3).

The loss of beta cells in type 1 diabetes (T1D) occurs over a period of several years, or even decades, both before and after clinical diagnosis. Insulin deficiency could tentatively be caused by 1) a specific loss of beta cells, the currently dominating view, or 2) by an inability to establish a beta-cell mass large enough to meet increased physiological demands. The relative importance of the two alternatives can be tested by morphometry of pancreases of subjects with or without T1D. We postulate that, if T1D is mainly caused by specific destruction of the beta cells, islets size would be reduced, since this cell-type constitutes about 65% of all cells within an islet (4), but the total number of islets would be preserved. Also, islets with signs of preceding beta-cell destruction, i.e. fibrosis, would be frequently found. Alternatively, if the main cause of insulin deficiency is an inability to establish islet mass during childhood and adolescence, a reduced number of islets would be found. In this study, we therefore compared the total islet number/mm^2^ pancreatic parenchyma (islet density) and the islet-size distribution in pancreatic biopsies obtained from patients with recent-onset T1D included in the Diabetes Virus Detection-study (DiViD) (5) and organ donors with T1D since several years, with that in matched non-diabetic organ donors (Table 1).

**Table 1.** Clinical data of patient cases with type 1 diabetes mellitus (T1D) and non-diabetic organ donors (Control donors).

| Variable | Control donors | Patients with recent onset T1D | Organ donors with longstanding T1D |
| --- | --- | --- | --- |
| Number of subjects | 9 | 6 | 7 |
| Age (years) | 25.7 ± 6.4 | 28.8 ± 5.1 | 36.6 ± 15.8 |
| Female | 3 | 3 | 3 |
| Male | 6 | 3 | 4 |
| Body mass index (kg/m<sup>2</sup>) | 25.4 ± 2.8 | 24.4 ± 3.1 | 23.9 ± 2.7 |
| HbA1c [% (mmol/mol)] | 5.6 ± 0.33<br>(38 ± 3.6) | 7.7 ± 1.3<br>(60 ± 14) | 8.2 ± 0.7<br>(10.5 ± 1.1) |
| Duration of T1D (weeks) | N/A | 5.2 ± 2.0 | > 200 |
| Donors with anti-GAD | 0 | 6 | 2 |
| Donors with anti-IA2 | 0 | 5 | 2 |
| Donors with anti-insulin | N/D | 3 | N/D |
| Donors with anti-ZnT8 | N/D | 3 | N/D |

Anti-GAD, anti-glutamic acid decarboxylase antibodies; anti-IA2, insulinoma-associated autoantibodies; Anti-ZnT8, zinc transporter autoantibodies. N/A, not applicable; N/D, not determined

## Research design and methods

### Biopsies

Pancreatic biopsies were obtained from six patients with recent-onset T1D, included in the DiViD study. Mean age was 28 years (range 24-35). Biopsies were taken laparoscopically from the pancreas tail 3-9 weeks after diagnosis. Two tissue samples from separate parts of each biopsy were analyzed, except in case 2 where three tissue samples were analyzed. Additional information regarding clinical history and surgery has been described previously (5). The DiViD study was approved by the Norwegian Governments Regional Ethics Committee and all patients gave informed written consent after having obtained oral and written information. Pancreatic tail biopsies were also collected from seven brain-dead organ donors enrolled into the Nordic Network for Islet Transplantation, with a history of more than 200 weeks (3.8 years) of type 1 diabetes, and from nine non-diabetic organ donors. The non-diabetic organ donors were age-matched to the DiViD patients with recent onset T1D (Table 1). Consent for organ donation (for use in research) was obtained via an online database (https://www.socialstyrelsen.se/donationsregistret/anmalan)or verbally from the deceased’s next of kin by the attending physician and documented in the medical records of the deceased in accordance with Swedish law and was approved by the Regional Ethics Committee (Dnr 2015/444). No diabetes-related autoantibodies were found in any of the control donors and glycated hemoglobin (HbA1c) levels were within the normal range (table 1).

### Sectioning and staining

Biopsies were fixed in formalin and embedded in paraffin. Biopsies from the DiViD collection were cut in 4 *µ*m sections in Oslo, Norway, and subsequently double-stained for glucagon (rabbit anti-glucagon antibody, clone EP74, dilution 1:400, Epitomics, Burlingame, CA, USA) and CD3 (rabbit anti-CD3 antibody, clone: SP7, dilution 1:150, Abcam, Cambridge, U.K.). Consecutive sections were double-stained for insulin (mouse anti-insulin antibody, clone 2D11/H5, dilution 1:100, Novocastra, Newcastle upon Tyne, U.K.) and CD3 (SP7). These sections were thereafter sent to Uppsala, Sweden for analysis. Sections in close proximity, but not directly consecutive, to the other sections were stained for synaptophysin (anti-synaptophysin clone DAK-SYNAP, dilution 1:100, Agilent, Santa Clara, CA, USA) in Uppsala. Biopsies from the donors in the control group were cut in 6 *µ*m sections and stained in Uppsala in the same manner as the sections from the DiViD samples. Biopsies from the donors with long standing T1D were double-stained for synaptophysin (DAK-SYNAP) and CD45 (mouse anti-human clone 2B11+PD7/26, dilution 1:75, Abcam), Proinsulin (mouse monoclonal, clone: C-PEP-01, dilution: 1:20000, ab7761 Abcam) and Glucagon (EP74), and CD3 (rabbit polyclonal, dilution 1:100, Agilent, cat:A0452) and Insulin (Guinea pig polyclonal anti-Insulin, dilution 1:200, Agilent, cat: A0564). Visualization was done using EnVision G|2 Doublestain System, Rabbit/Mouse (Agilent, Cat: K536111-2)

All slides were scanned with the Aperio ScanScope system (Aperio Technologies, Oxford, UK). In the sections stained for synaptophysin, the borders of each pancreatic islet were identified morphologically and marked using the pen tool in Aperio ImageScope. The total tissue area in each section was calculated using the pen tool and the negative pen tool. Morphological identification of islets using the pen tool was also applied on the sections stained for glucagon/CD3 and insulin/CD3. The glucagon- and insulin-containing areas were calculated in the sections from the DiViD patients and from the non-diabetic control donors, using the Positive Pixel Count Algorithm in Aperio ImageScope version 12.3 software (Aperio Technologies). All islets with an area >491 *µ*m^2^ (corresponding to a diameter of >25 *µ*m), approximately corresponding to islets containing a minimum of five glucagon- or insulin-containing cells, were regarded for analysis. Circularity is a measure of the roundness of an object, it can be calculated when the area and perimeter is known through the formula: C = 4πA/P^2^, where C is the circularity, A is the area and P is the perimeter. A perfect circle has C = 1. In total, 12.7 cm^2^ pancreatic tissue from patients with recent onset T1D, 6.81 cm^2^ from donors with longstanding T1D, and 4.97 cm^2^ from control organ donors was analyzed.

### Multiplex staining and analysis

The Opal 7-Color Automation IHC Kit (Perkin Elmer, Waltham, MA, USA, Cat: NEL801001KT) was used for multiplex staining as per manufacturer’s instructions. A series of regular immunohistochemical stainings was performed, each separated by microwave heating to strip the tissue of previously bound antibody. Opal fluorophores bind covalently to the antigen and remain bound after microwave treatment. In brief, paraffin sections were deparaffinized in xylene, rehydrated in alcohol, and fixated in 4% PFA for 20 minutes. Blocking was performed using Antibody Diluent/Block for 10 minutes at RT. Stainings were done in the following order: Insulin (guinea-pig polyclonal-anti insulin, dilution 1:100, Agilent, Cat: A0564), DAKO Envision+ HRP labelled anti-rabbit polymer (Agilent, Cat: K4003). Glucagon (rabbit anti-glucagon antibody, clone EP74, dilution 1:400, Epitomics, USA), DAKO Envision+ HRP labelled anti-rabbit polymer (DAKO, Cat: K4003). PDX-1 (polyclonal goat anti-PDX1, 15ug/mL, R&D Systems, Minneapolis, MN, USA, Cat: AF2419), Anti-goat (Donkey anti-goat, dilution 1:2000, Abcam, Cat: ab6885). ARX (polyclonal sheep anti-ARX, 20ug/mL, R&D Systems, Cat: AF7068). Anti-sheep (Donkey anti-sheep, dilution 1:2000, Abcam, Cat: ab6900) Visualization was done with Opal 520, Opal 570, Opal 540 and Opal 620, respectively. Sections were counterstained with Spectral DAPI. In between each IHC series, sections were heated in a microwave in citrate buffer pH 6 for 1 minute at 1000W, then 15 minutes at 100W. Images were taken with the Vectra Polaris platform (Perkin Elmer) at 20X magnification and the fluorescent signal was unmixed using inForm 2.42 software (Perkin Elmer).

### Data analysis

Calculations and statistical analyses were carried out using R version 3.3.0. Differences between groups were analyzed using non-parametric ANOVA (Kruskal-Wallis) and the Mann-Whitney U-test.

## Results

### Morphological characterization of islets in non-diabetic subjects, subjects with recent onset or longstanding T1D

Islets with various morphological appearance were seen in patients with recent onset T1D. In 97% (92-100%) of the islets in non-diabetic subjects, ≥60% of the endocrine area stained positive for insulin. In subjects with recent onset T1D, 13% (3.0-40%) of the islets contained ≥60% insulin, 13% (5.1-22%) contained 20-60% insulin and 74% (47-90%) contained ≤20% insulin (fig 1). In subjects with longstanding T1D, no insulin-containing islets were found and the absolute majority of the islet cells expressed glucagon. However, in 5 of the 7 subjects with longstanding T1D, single insulin-positive cells were found scattered throughout the exocrine parenchyma, albeit to a lower extent than in controls.

**Figure 1:**
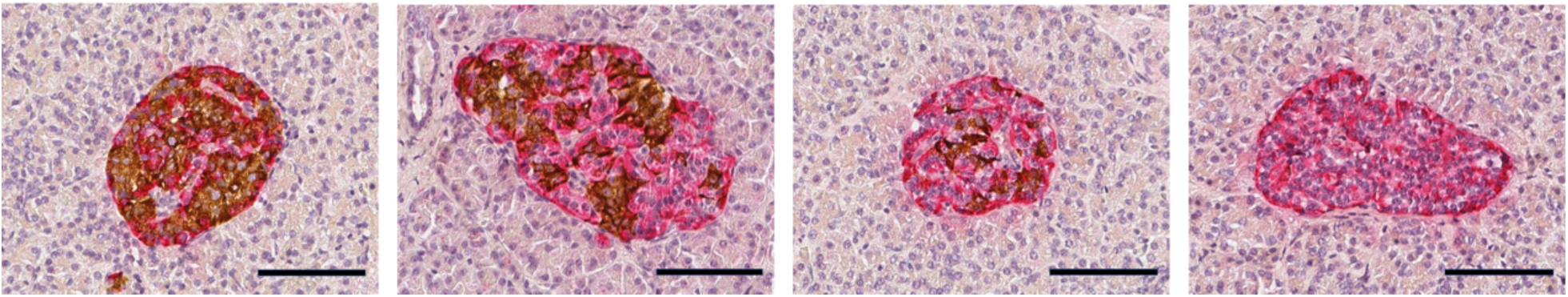
Examples of islets with reduced insulin content and preserved architecture in subjects with recent-onset type 1 diabetes. Islets stained for insulin (brown) and glucagon (red). Insulin-deficient islets seem to be preserved in size and the absolute majority of cells express glucagon. No necrotic or fibrotic areas were noted in the islets. Scale bars 100 um.

Almost all islets were seemingly intact without discernable necrotic or large fibrotic areas regardless of disease status. Some islets having irregular borders were observed, especially among subjects with recent-onset T1D but most islets had a normal architecture. Circularity was used to measure this phenomenon and, although there was a tendency to lower circularity when comparing recent-onset T1D subjects with control subjects, no statistically significant differences were seen (fig 2a).

**Figure 2:**
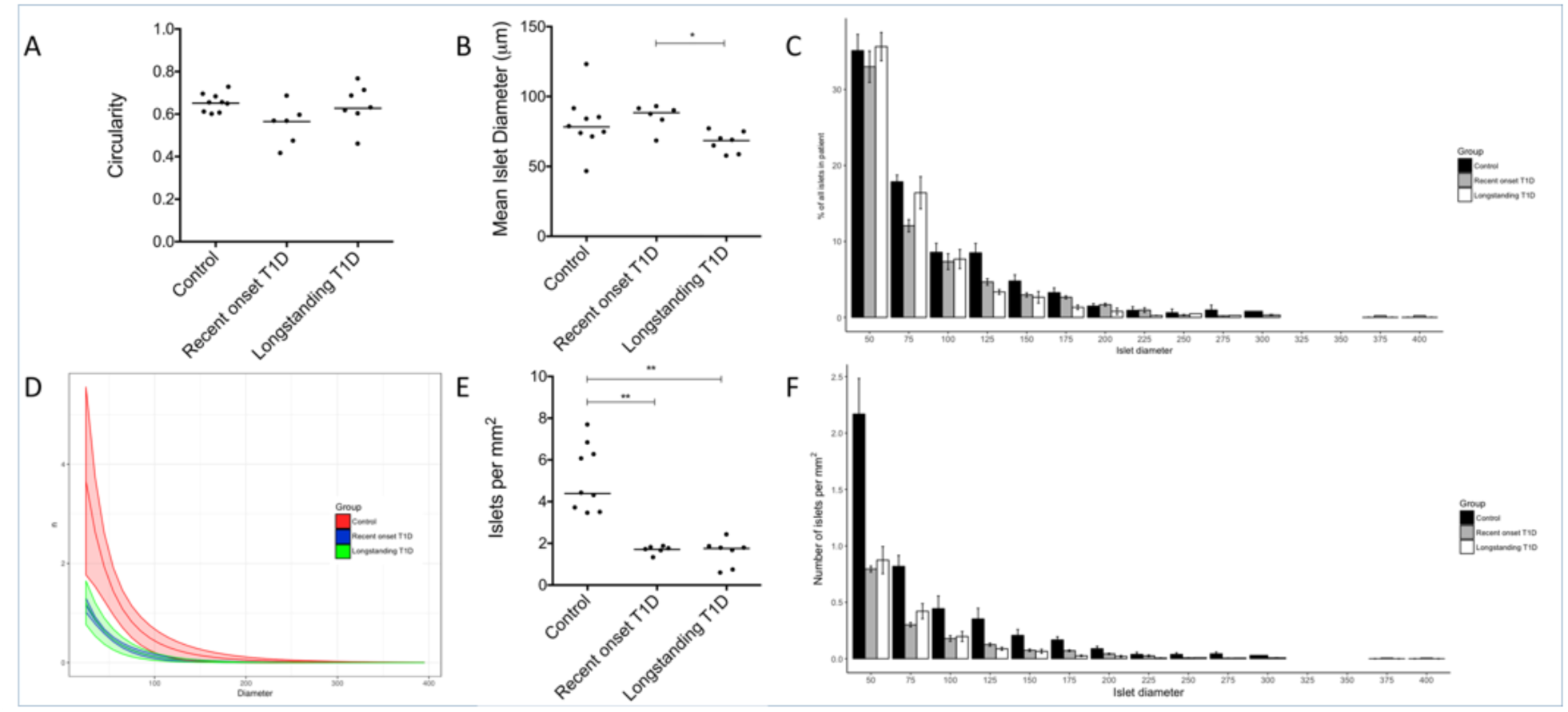
Comparison between patients with recent-onset T1D and organ donors without diabetes. (A) Mean circularity of islets in each donor (dots) and the median for each donor group. Circularity is calculated by the formula C = 4πA/P^2^, where C is the circularity, A is the area and P is the perimeter. A perfect circle has C = 1. (B) Mean islet diameter in each donor (dots) and the median for each donor group (horizontal line). (C) Percentages of islets in each size category in steps of 25 um in islet diameter. (D) The mean exponential curve (y = a × e ^−bx^) fitted to the mean number of islets per mm^2^ in each size category for each donor. (E) Mean islets/mm^2^ in each donor (dots) and the median for each donor group (horizontal line). (F) Number of islets per mm^2^ in each size category in steps of 25 um in islet diameter. * denotes a significance of <0.05. ** denotes a significance of <0.005.

### Islet size distribution

The percentage of islets in each size category was calculated in relation to the total number of islets (fig 2c). There was no difference in islet size distribution between the three groups of subjects. This was confirmed by fitting an exponential curve (y = a × e ^−bx^) to the mean number of islets per mm^2^ per islet size category in each donor (fig 2d). No statistically significant differences were seen between any of the groups comparing the values for the slope coefficient *b* between the groups (*b* in the control group was 0.028, 0.030 in recent onset T1D and 0.034 in longstanding T1D).

The median of the mean islet size in each donor was 4813 (1673-11807) *µ*m^2^ in controls, as compared to 6127 (3623-6721) *µ*m^2^ in patients with recent onset T1D, and 3678 (2569-4608) *µ*m^2^ in donors with longstanding T1D, corresponding to a diameter of 78 (46-123) *µ*m in controls, 88 (68-93) *µ*m in recent onset, and 68 (57-77) *µ*m in longstanding T1D (fig 2b). The differences in mean islet size between controls and either of the groups of subjects with T1D were not statistically significant. However, islets from subjects with recent onset T1D was significantly larger than from subjects with longstanding T1D (p<0.05).

### Islet density in the pancreas

A total of 2054, 1075, and 2197 islets were counted in subjects with recent onset T1D, longstanding T1D, and non-diabetic controls, respectively. The median islet density was significantly lower in subjects with recent onset T1D (1.7 islets/mm^2^, range 1.3-1.8) or longstanding T1D (1.8 islets/mm^2^, range 0.6-2.4) compared to non-diabetic control subjects (4.4 islets/mm2, range 3.4-7.7) (p<0.005). No difference in islet density was found between subjects with recent onset T1D and subjects with longstanding T1D (fig 2e).

The number of islets/mm^2^ pancreas in each size category was calculated (fig 2f). In each size category the number of islets/mm^2^ was reduced with about 50% in both categories of diabetic subjects when compared to non-diabetic subjects. This was confirmed in the exponential curve described above (fig 2d), and then comparing the values for the constant *a* between the groups. The mean *a*-value in the control group was 9.0, significantly (p<0.0115) higher than in the two T1D groups (2.5 in recent onset and 2.8 in longstanding T1D). No statistically significant differences were seen between subjects with recent onset and longstanding T1D.

### Altered expression of transcription factors in alpha-cells in recent-onset and long-standing type 1 diabetes

In islets from non-diabetic subjects, and in insulin-containing islets with apparent normal number of insulin-containing cells from the DIVID subjects, the beta-cell transcription factor PDX1 was localized in nuclei of insulin-containing cells, and the alpha-cell transcription factor ARX localized in nuclei of glucagon-containing cells, as expected. However, PDX1 expression was frequently found in nuclei of glucagon-containing cells in islets with reduced number of insulin-containing cells in recent-onset subjects and in islets without insulin-containing cells in both recent-onset and long-standing T1D subjects (fig 3). No insulin-containing cell expressing nuclear staining with ARX was found and no cells were found with discernable co-expression of glucagon and insulin.

**Figure 3:**
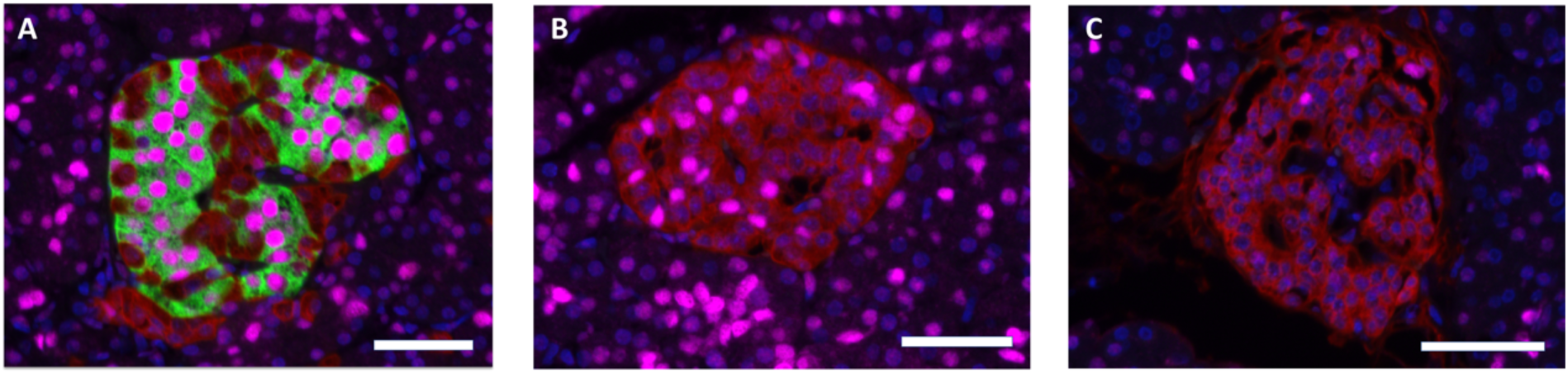
PDX1-expression in alpha cells of recent-onset and longstanding type 1 diabetic subjects. Sections stained for insulin (green), glucagon (red), PDX1 (magenta). Scale bars 50 um. (A) Insulin-containing islet in a recent-onset T1D subject displaying PDX1 expression in beta-cells but not in glucagon-positive cells. (B) Insulin-deficient islet from the same pancreatic section as (A) displaying PDX1 expression in alpha-cells. This islet is situated only ~500 um from the islet in A. (C) Islet from a subject with long-standing T1D, negative for insulin but with a few alpha-cells expressing PDX1.

## Discussion

The results presented show preserved islet size distribution, but a marked reduction in islet density in subjects with recent onset T1D when compared with non-diabetic subjects and that no further reduction in islet numbers/mm^2^ occurs with increased disease duration. Several previous publications have shown a marked reduction in beta cell volume in T1D; at diagnosis only about 1/3 or less of the presumed beta cell volume remains (6). A so far neglected area of investigation has been the actual number of islets present in subjects with T1D; in non-diabetic subjects islet density averages about 25 islets/mm^2^ during childhood and is less than 10 islets/mm^2^ in adults (2). Only few studies have indirectly reported data from which the number of islets per volume or area of pancreas can be extracted. Importantly, these studies align with the herein presented results, i.e. a reduction in the range of 30-70% in subjects affected by T1D when compared to non-diabetic controls (7-13).

Our observations of reduced number of islets/mm^2^ in T1D should be viewed in relation to the apparent absence in the literature of reports on significant beta-cell destruction or apoptosis in subjects with T1D (14). Almost all publications in humans affected by T1D show few T cells residing in the peri-islet area of only few islets but without aggressive T cell infiltration and ongoing beta cell destruction. Also, no correlation has been found between remaining beta cell mass and the frequency of insulitis (15).

The islet size distribution as well as the mean islet size was similar in all three groups examined herein. These results indicate that the remaining islets in both subjects with recent onset as well as longstanding T1D contain a similar number of cells, but with a markedly changed phenotype of cells. In subjects with longstanding T1D, almost all remaining cells within the islets were alpha cells. Also, most islets in the DiViD subjects contained significantly larger proportion of alpha cells when compared with that in non-diabetic subjects. Already in 1978, Gepts et al reported on increased relative numbers of alpha-cells in both recent-onset and long standing T1D patients (8). An important role for glucagon in T1D pathology has been suggested and exaggerated plasma glucagon responses to mixed-meal are observed in children and adolescents with T1D within the first 2 years of diagnosis (16).

Focal lesions of acute pancreatitis with accumulation of leukocytes, often located around the ducts, are frequently observed in subjects with recent onset T1D (8) and most patients display extensive periductal fibrosis, the end stage of inflammation (17). An injurious inflammatory adverse event occurring within the periductal area may have negative implications for islet neogenesis; dependent on stem cells residing within or adjacent to the ductal epithelium (3; 18). Failure to expand beta-cell mass during childhood would lead to prolonged periods of beta cell exhaustion and finally clinically overt T1D. Several studies in subjects with recent-onset T1D have shown an almost absent insulin-secretory response after intravenous injection of glucose. However, in response to oral glucose (triggering the incretin effect) or after an intravenous injection of arginine, these subjects release significant amounts of insulin (19; 20). This “glucose-blindness” can conceivably be considered as the first sign of beta cell exhaustion.

The first report on dedifferentiation of beta cells in rats with chronic hyperglycemia was published already in 1999 (21). The transcription factor FOXO1 has key role in the process: lineage-tracing experiments in mice with somatic deletion of FOXO1 in beta cells showed reduced numbers of beta cells caused by dedifferentiation to alpha cells, not beta cell death, in situations with beta cell stress (22). These results were recently corroborated also in human type 2 diabetes (T2D) were a large fraction of beta cells dedifferentiated to alpha or delta cell phenotypes (23).

Notably, dissociation of human islets into single cells, and subsequent reaggregation into clusters, cause degranulation of beta cells that seems to trigger dedifferentiation of most beta cells into glucagon expressing cells (24) but with retained expression of the beta cell specific transcription factors PDX1 and NKX6.1.

Surprisingly, this process of beta cell loss has not so far been explored in T1D. Similar to the herein reported finding of PDX1 reactivity in the nuclei in a large proportion of the alpha cells, a recent publication on impaired alpha cell function in T1D showed that most alpha cells in donors with T1D do not express the alpha cell transcription factors MAFB and ARX, but low levels of NKX6.1 (25). Obtained findings were interpreted as that only a partial change from an alpha toward a beta cell phenotype had occurred. Hence, similar observations in T1D and T2D have been interpreted in different ways: in T2D to support the hypothesis of beta cell dedifferentiation to alpha cells as a mechanism for beta cell loss, whereas in T1D to support the hypothesis of beta cell neogenesis from alpha cells.

Assessment of the total number of islets within the pancreas would have required access to the entire pancreas, however, only the tail of the pancreas was resected in the DiViD subjects. Islet density in non-diabetic subjects has been reported to be slightly increased (≈10%) in the tail when compared to the head and corpus and the results presented here may not be representative for the whole pancreas. The results should, however, be comparable between all groups as the tail was examined also in the organ donor pancreases.

Collectively, our findings suggest that acute massive beta-cell destruction in T1D is not occurring recently after clinical diagnosis. Neither has this been described in other published reports on subjects with recent onset T1D. We suggest that dedifferentiation of beta cells could explain a maintained islet size distribution and that loss of endocrine mass is mainly due to processes affecting entire islets.

## Acknowledgements

The authors thank specialist nurse T. Roald, Oslo University Hospital, Norway, who provided invaluable efforts in coordinating the study; and to the surgeons Professor Bjørn Edwin and Professor Trond Buanæs at Oslo University Hospital providing the tissue biopsies in the DiViD Study. We thank also nurses and doctors at the local hospitals providing contact with the patients, Sofie Ingvast for skillful technical assistance, and the patients who participated in this study.

## Author contributions

A.W. and P.S. performed the data analysis, interpreted the data and drafted the manuscript. L.K. was responsible for clinical coordination and patient recruitment within the DiViD study. KD-J was the principal investigator of the DiViD study and is responsible for the study design, funding, regulatory issues, and international collaboration of the DiViD study. FLJ performed the insulin and glucagon staining of DiViD pancreatic biopsies. O.S. and O.K. planned the study and the study design, interpretation of the results, contributed to discussion and writing of the manuscript. O.K. is the guarantor of this work and, as such, had full access to all the data in the study and takes responsibility for the integrity of the data and the accuracy of the data analysis. All authors had final approval of the version to be published.

